# Diffusion MRI based biomarkers reveal a Prolonged Pre-Lesional Phase of Cerebral Small Vessel Disease

**DOI:** 10.64898/2026.03.03.709452

**Authors:** Prashanthi Vemuri, Mingzhao Hu, Emily S. Lundt, Michael G. Kamykowski, Robert I. Reid, Terry M. Therneau, Sheelakumari Raghavan, Petrice M. Cogswell, Michael Griswold, B. Gwen Windham, Clifford R. Jack, Ronald C. Petersen, Jonathan Graff-Radford

**Affiliations:** Department of Radiology, Mayo Clinic-Rochester, Rochester, Minnesota, USA; Department of Quantitative Health Sciences, Division of Clinical Trials and Biostatistics, Mayo Clinic-Rochester, Rochester, Minnesota, USA; Department of Information Technology, Mayo Clinic-Rochester, Rochester, Minnesota, USA; The MIND Center, University of Mississippi Medical Center, Jackson, Mississippi, USA; Department of Neurology, Mayo Clinic-Rochester, Rochester, Minnesota, USA

**Keywords:** small vessel disease, WMH, temporal progression, diffusion MRI

## Abstract

**Background:** White matter hyperintensities (WMH) are widely used to assess cerebral small vessel disease (SVD) but reflect late-stage injury. Diffusion MRI based biomarkers have been proposed to capture earlier SVD-related microstructural damage but their temporal progression relative to WMH and the risk factors associated with this progression have not been explored.

**Methods:** We analyzed longitudinal neuroimaging data from 2,047 participants from a population-based cohort study (aged 49-101 years, 47% female). Using multi-output nonlinear mixed-effects models, we characterized the temporal progression of WMH and four diffusion MRI based biomarkers: fractional anisotropy of the genu of the corpus callosum (Genu-FA), peak width of skeletonized mean diffusivity (PSMD), free water (FW), and Arteriolosclerosis-score (ARTS). Models incorporated participant-specific time shifts, correlations between biomarkers, and effects of risk factors (sex, education, *APOE ε4* status, and cardiometabolic conditions).

**Results:** ARTS, Genu-FA, FW, and PSMD became abnormal in 50% of the study population 16, 12, 10, and 7 years before WMH, respectively. Global markers (ARTS, FW, PSMD, WMH) were correlated, indicating shared substrates of widespread white matter injury. Genu-FA, a vascular risk microstructural injury biomarker, was weakly coupled with WMH and had an earlier but more linear worsening across adulthood. Cardiometabolic conditions predicted earlier worsening of all biomarkers. Females showed earlier WMH, Genu-FA, and ARTS abnormalities whereas males exhibited earlier PSMD and FW abnormalities.

**Conclusions:** Diffusion MRI based biomarkers capture microstructural injury at least a decade before appearance of WMH, revealing a prolonged phase of early SVD and highlighting their potential for SVD prevention.

## BACKGROUND

Small vessel disease (SVD) is associated with poor cognitive, gait, and brain health outcomes^1^ and is primarily driven through the mechanisms of arteriolosclerosis (associated with hypertension) and cerebral amyloid angiopathy^2^. Worsening SVD manifests as deteriorating white matter (WM) health, owing to the distinctive vulnerabilities of WM vasculature and metabolism^3^. Further, poor WM health is also seen with dying back (degeneration) of connectivity in the presence of cerebrovascular lesions^4^. While white matter hyperintensities (WMH) are traditionally clinically utilized for assessment of SVD, there is increasing evidence that early subtle changes in WM can be quantified using fractional anisotropy (FA) and mean diffusivity (MD) measures from diffusion MRI and reflect early SVD damage^5-7^. However, there has been limited effort to understand how early these recently proposed SVD biomarkers change relative to WMH. Understanding this important knowledge gap will aid in accelerating early prevention and treatment strategies for SVD.

Early SVD can be measured using regional or global WM microstructural injury alone or in combination with other features^8^. Fractional anisotropy of the genu of the corpus callosum (frontal interhemispheric WM; Genu-FA) has demonstrated heightened sensitivity to vascular risk factors, as shown by our group and others^9,10^, potentially due to the region’s structural fragility and relatively poor collateral vascular supply. Peak width of skeletonized mean diffusivity (PSMD)^11^ and free water (FW)^12^ reflect global WM damage and have been proposed as key SVD biomarkers in the MarkVCID consortium^13^ along with the Arteriolosclerosis-score (ARTS)^14^. ARTS provides *in vivo* measurement of arteriolosclerosis combining regional diffusion MRI features with WMH and demographics. Given the need to understand how these biomarkers evolve overtime relative to one another, temporal modeling approaches such as multi-output nonlinear mixed effects (NLME) that have been effective for Alzheimer’s disease biomarker modeling are well suited, as they can provide a natural framework for SVD progression^15,16^. NLME models provide smooth, monotone progression curves for multiple biomarkers on a shared age axis, with individual-level time shifts indicating whether a given participant progressed earlier or later than average for each marker relative to the population mean.

The primary goal of this paper was to model the temporal progression of the above-mentioned four early SVD biomarkers (Genu-FA, PSMD, FW, ARTS) relative to WMH in a population-based sample where the complete range of SVD is observed in participants who have extensive longitudinal imaging data. A secondary goal was to evaluate the impact of demographics, vascular risk factors, and *APOE ε4* carriership on these longitudinal trajectories to identify risk profiles for these biomarkers.

## METHODS

### Participants

Participants were drawn from the Mayo Clinic Study of Aging (MCSA), an ongoing, population-based longitudinal cohort based in Olmsted County, Minnesota^17-20^. Recruitment for the MCSA began in 2004 with individuals aged 70 to 89 years. In subsequent expansions, eligibility was broadened to include adults aged 50 and older in 2012, and further to individuals aged 30 and above in 2015. For this study, we included MCSA participants who were aged 50 or older and had at least two imaging visits for at least one imaging SVD biomarker to aid longitudinal modeling. Demographic variables, including age, sex, and years of education, were collected as part of the baseline clinical evaluation. *APOE ε4* genotype was available in all study participants. Clinical diagnosis of cognitively unimpaired (CU), mild cognitive impairment (MCI), or dementia was ascertained at each visit through consensus conferences. The presence and absence of seven cardiovascular metabolic conditions (hypertension, diabetes, dyslipidemia, stroke, congestive heart failure, coronary heart disease, cardiac arrhythmias) based on nurse abstraction from medical records were summarized as the cardiovascular metabolic conditions (CMC) score^21^. As of September 2024, 7,437 MCSA participants had at least one study visit. After excluding participants missing baseline covariates required for modeling (n = 1,127, with overlap across covariates), 6,310 remained. Of these, 2,802 were excluded for not having an eligible baseline MRI biomarker measurement, 220 were excluded for not meeting age 50+ or clinical diagnosis criteria at the most recent visit, and 1,241 were excluded for not having a follow up biomarker measurement. The final sample included 2,047 for analysis.

All available diffusion MRI and FLAIR-MRI observations were included, and participants contributed information only at observed visits. Missing outcome data was not imputed; inference was likelihood based under an assumption of Missing at Random. Participants with missing baseline characteristics (age, sex, education, CMC, *APOE ε4* genotype, clinical diagnosis) were excluded prior to analysis. Participants were required to have at least two observed biomarker measurements to allow estimation of longitudinal trajectories.

### Standard protocol approvals, registrations, and patient consents

The study was approved by the Mayo Clinic and Olmsted Medical Center institutional review boards and was performed in accordance with the ethical standards of the Declaration of Helsinki and its later amendments. All participants or a legally authorized representative provided informed written consent.

### MRI acquisition and processing

Magnetic resonance imaging (MRI) was performed on 3 Tesla scanners (GE Healthcare and Siemens). The imaging sequences utilized for this work included T1-weighted MPRAGE, FLAIR, and diffusion MRI. An automated method was used for computing WMH (which is scaled by the total intracranial volume)^22^ and regional FA measures (for Genu-FA) were computed using JHU atlas as previously reported^8^. We computed the FW and arteriolosclerosis (ARTS)-score using MarkVCID scripts^13^, while PSMD was calculated using a freely available fully automated script (v 1.8.3 -http://www.psmd-marker.com/), as previously published^11^. In the MCSA, a platform transition from GE Medical Systems scanners to Siemens Prisma scanners occurred in Fall 2017. We utilized deming regression approach from data acquired on 111 participants on both GE and Siemens scanners within a week to harmonize the data on both vendor platforms^23^.

### Primary model and individual adjustments

We modeled biomarker progression using a NLME framework, also previously described as an accelerated failure time (AFT) approach and used for Alzheimer’s disease^15,16^. The outcomes included one macrostructural biomarker (WMH) and four diffusion MRI based biomarkers: Genu-FA, FW, ARTS, PSMD. WMH was transformed using a square root function to correct for skewness. Genu-FA was scaled as (1−Genu-FA)×5−1, PSMD as log(PSMD)×2+18, FW as FW×8−1, and ARTS as ARTS×1.2+1.2. This transformation prior to model fitting was to enforce monotonic increases over age and to standardize the scale across outcomes, thereby improving numerical stability and reducing convergence time. All model results were transformed back to native units for interpretation and reporting. The model specification included three primary components: (1) a non-linear progression function for each biomarker over age, chosen to be the simplest monotonic form sufficient to capture temporal trends; (2) fixed effects linking covariates to each outcome; and (3) a subject-specific random effect for each biomarker, reflecting left–right individual-level variation in onset age, based on baseline covariates. These individual adjustments can also be viewed as time shifts per participant trajectory over age. As part of the model output, we obtained a correlation matrix quantifying the relationships among random effects, representing the strength of association in individual-level age adjustments between each biomarker pair. Covariate coefficients were thus estimated as age shifts (in years) in the timing of each biomarker’s trajectory. Additional technical details on model implementation and validation are available in prior work^15,16^. Loss to follow up was handled implicitly by the likelihood based NLME using all observed measurements, without imputation. This approach is valid under a Missing at Random assumption for dropout conditional on observed history and included covariates. We did not fit a separate model for informative dropout.

### Cut-point determination and population-level diffusion marker progression

To define biomarker abnormality, we applied empirically derived thresholds validated in the MCSA. For WMH, a square-root-transformed value of 0.98 was chosen, corresponding to a Fazekas-equivalent cutoff of ≥2 indicative of moderate to severe WMH burden^22^. For the diffusion MRI metrics, cut-points were established using a data-driven method designed to maximize both sensitivity and positive predictive values comparing individuals with vascular cognitive impairment (based on WMH and clinical impairment) with younger (30–49-year-old) cognitively unimpaired participants. The resulting thresholds for abnormality were: 0.37 for 1 – Genu-FA (equivalent to less than 0.63 for Genu-FA), 0.00017 for PSMD, 0.22 for FW, and -0.47 for ARTS. These cut-points were applied in downstream analyses to characterize the age-related trajectory of abnormal biomarker expression at the population level. More details on the framework we utilized to select the best cut-point can be found in Hu et al.^23^.

## RESULTS

### Participants

The sample included participants with a mean age of 78 years, of whom 53% were male and 28% *APOE ε4* allele carriers (**Table 1**). At their most recent clinical assessment, 81% were classified as CU, while 17% were diagnosed with MCI, and 3% met criteria for dementia. The number of participants with at least two time point measurements for each biomarker were WMH (98%), FA (62%), PSMD (64%), FW (63%), and ARTS (64%). The lower percentage of diffusion MRI measures was because diffusion MRI started later in the MCSA. The average duration of imaging follow-up was over four years for all biomarkers, with WMH having the longest mean follow-up of 5.2 years (SD = 3.5), and other measures including Genu-FA, PSMD, FW, and ARTS averaging 4.2 to 4.3 years (SD = 3.9 for all).

**Table 1.** Participant characteristics. Statistics are mean (SD) or n (%) unless otherwise specified.

|  | Overall (N=2047) |
| --- | --- |
| Age at most recent visit, years | 78 (10) |
| Male | 1091 (53%) |
| Education, years | 15 (3) |
| APOE ε4 carrier | 564 (28%) |
| Diagnosis at most recent visit<sup>1</sup> |  |
| CU | 1651 (81%) |
| MCI | 338 (17%) |
| Dementia | 58 (3%) |
| Total no. FLAIR-MRI | 3 (1) |
| Total no. DTI-MRI | 2 (1) |
| Years of FLAIR-MRI follow-up | 5.2 (3.5) |
| Years of DTI-MRI follow-up | 4.3 (3.9) |
| Measurements at most recent visit |  |
| WMH | 1.060 (1.027) |
| Genu-FA | 0.629 (0.040) |
| PSMD | 0.00019 (0.00004) |
| FW | 0.234 (0.043) |
| ARTS | -0.325 (0.209) |

CU=cognitively unimpaired; MCI=mild cognitive impairment

### Model fits

Figure 1. displays the modeled trajectories of SVD biomarkers plotted against both chronological age and adjusted age. Adjusted age represents the covariate- and random effect–based estimate of a participant’s effective age relative to the onset of each biomarker’s trajectory. To ensure robust parameter estimation, the model was implemented in RSTAN with 4000 sampling chains. Convergence diagnostics were evaluated using the ratio of the between-chain variance to the within-chain variance (i.e., R-hat statistic), which approached 1.0 for all parameters, indicating proper mixing and convergence of the Markov chains. This was corroborated by large effective sample sizes, confirming that the posterior draws behaved as a sufficient set of independent samples.

**Figure 1:**
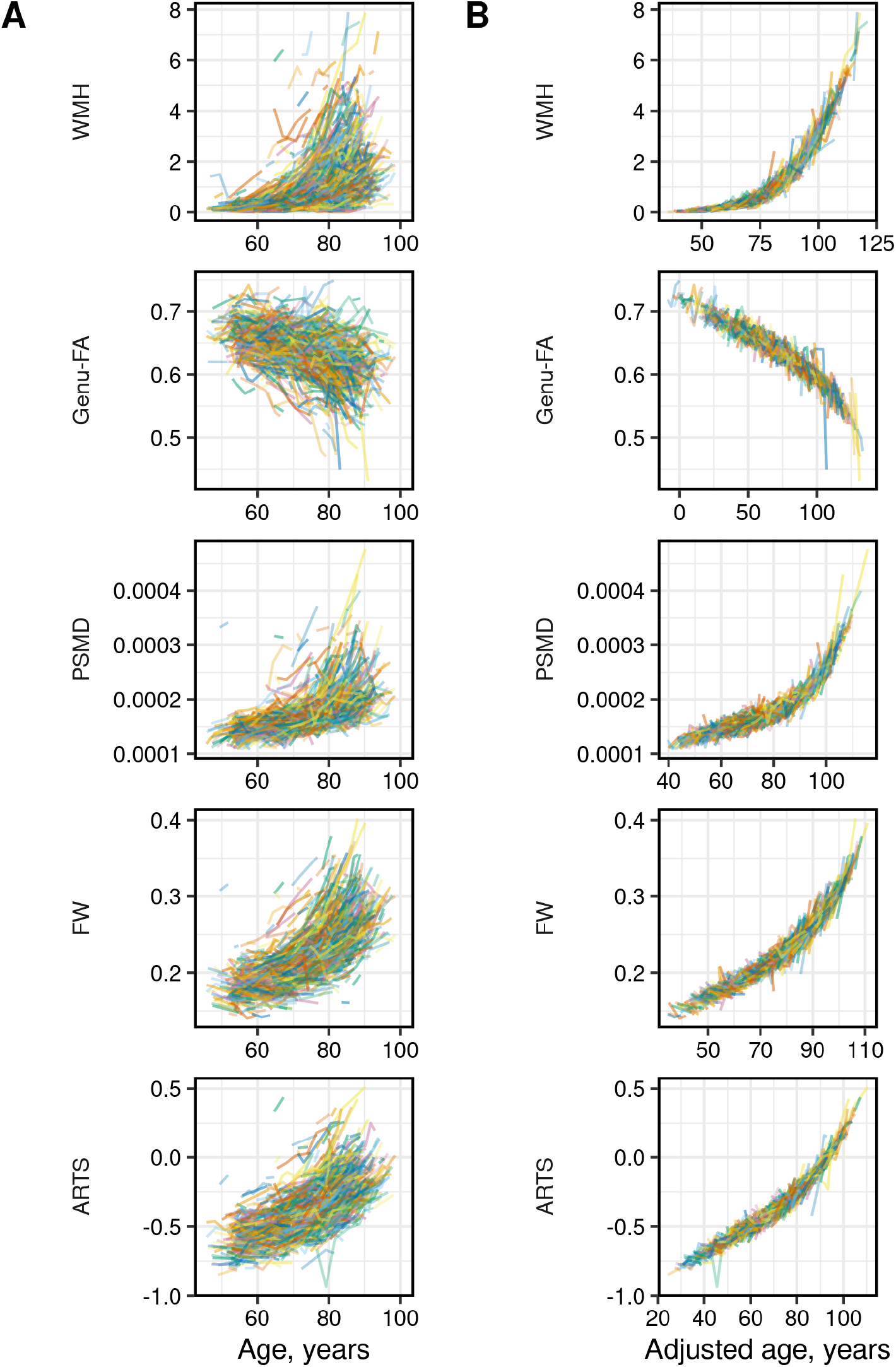
Trajectories of SVD biomarkers vs age (left panel), adjusted age (right panel). A scatter in color of trajectories from all participants to evaluate against the group mean curve to review the shape of the relationship. The trajectories of participants have been shifted left/right along the time scale in the adjusted age panel to correctly reflect this adjustment.

### Association of SVD biomarker timings

**Figure 2**. and **Table 2** summarize the correlations among participant-specific time shifts across SVD biomarkers. These adjustments reflect left–right shifts along the age axis for each biomarker trajectory. Strong associations were observed between WMH and early SVD biomarkers. The estimated correlation between WMH and PSMD adjustments was R = 0.75, with a 95% credible interval of (0.72, 0.77). Similarly, the individual shift in WMH timing showed moderate to strong correlation with that of FW (R = 0.60, 95% CI: 0.57, 0.64) and ARTS (R = 0.70, 95% CI: 0.67, 0.72) (**Table 2**). In contrast, the association between Genu-FA and WMH adjustments was weaker, with R = 0.39 (95% CI: 0.35, 0.44). Among the early SVD biomarkers themselves, the strongest correlations were observed between FW and PSMD (R = 0.82, 95% CI: 0.80, 0.84) and between FW and ARTS (R = 0.81, 95% CI: 0.79, 0.83). Interpretation of **Figure 2** is facilitated by considering these adjustments as per-person deviations from the mean age of biomarker accumulation. For example, in the WMH versus PSMD comparison, the 10th to 90th percentile range of individual WMH shift values spanned approximately –13 to +14 years, while PSMD ranged from about –11 to +12 years. For participants having approximately 20 years earlier than the mean age of onset for PSMD, the mean age of onset for WMH will be between 13 to 27 years earlier.

**Table 2.** Correlation coefficients, R (95% credible interval), between individual-level adjustments. The 100 x R^2^ values can be seen in Figure 2.

|  | WMH | Genu-FA | PSMD | FW |
| --- | --- | --- | --- | --- |
| Genu-FA | 0.39 (0.35, 0.44) |  |  |  |
| PSMD | 0.75 (0.72, 0.77) | 0.53 (0.49, 0.57) |  |  |
| FW | 0.60 (0.57, 0.64) | 0.69 (0.66, 0.72) | 0.82 (0.80, 0.84) |  |
| ARTS | 0.70 (0.67, 0.72) | 0.60 (0.57, 0.64) | 0.68 (0.65, 0.71) | 0.81 (0.79, 0.83) |

**Figure 2:**
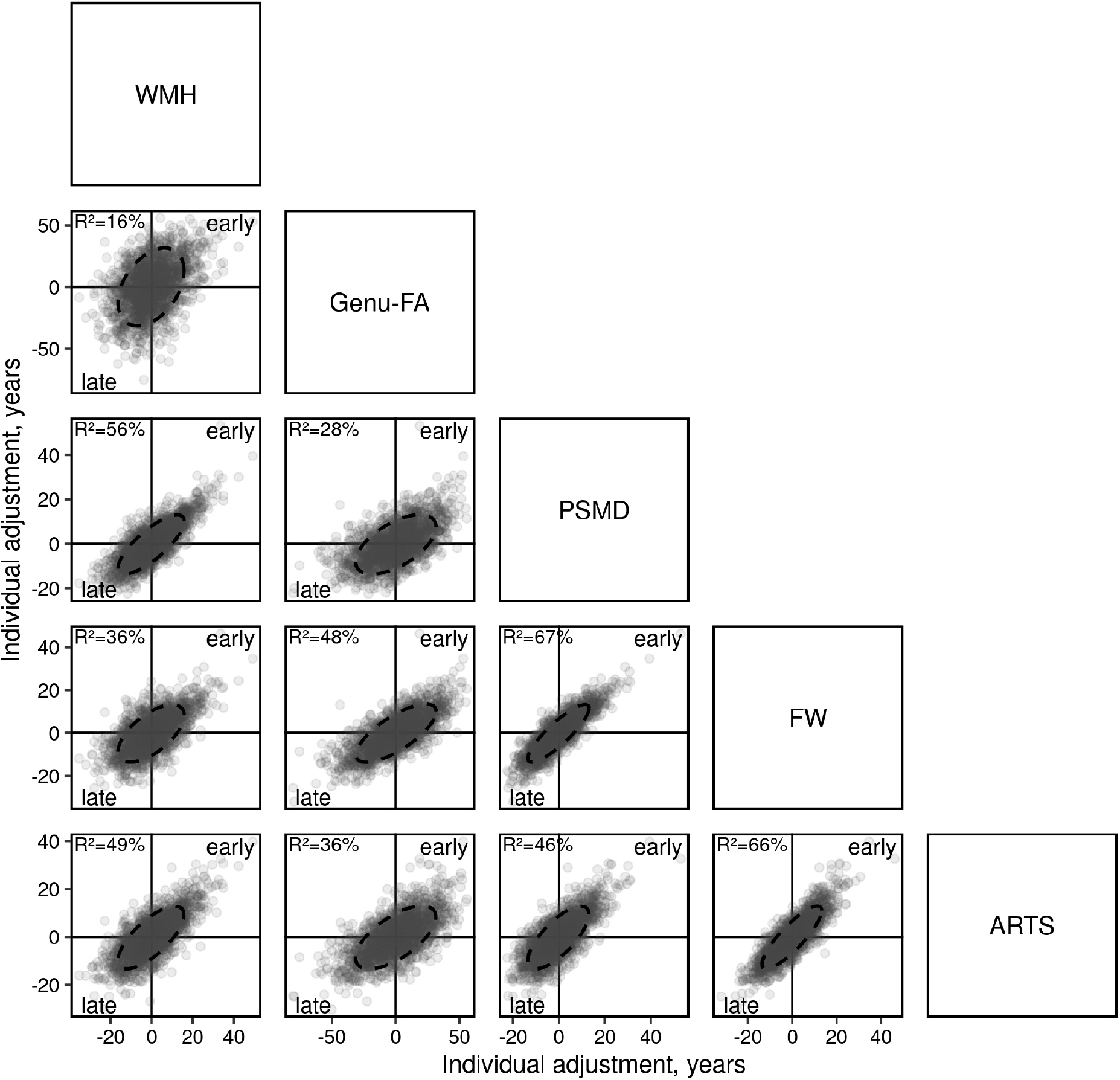
Correlation structure of participant-level time shifts across cerebrovascular biomarkers. Scatterplots display pairwise relationships between individual age adjustments for WMH, Genu-FA, PSMD, FW, and ARTS score. Each point corresponds to one participant, with the sample size varying across panels depending on biomarker data availability. Axes represent estimated deviations (in years) from the mean age of onset for each biomarker, with higher values indicating earlier-than-average onset (rightward along the x-axis or upward along the y-axis). Ellipses capture 80% of the data distribution; circular ellipses suggest minimal correlation between time shifts. The percentage of variance shared between paired adjustments (R^2^ × 100) is shown in the upper left corner of each plot.

### Covariate effects

Covariate effects on biomarker timing, expressed as left–right shifts in the age axis, are presented in **Figure 3** and **Table 3**. Among participants in the MCSA, sex showed the most pronounced association across SVD biomarkers, followed in strength by the burden of CMC. For WMH, Genu-FA, and ARTS, being male was associated with a delayed onset of abnormality by 2.8, 4.9, and 9.3 years, respectively. In contrast, being male was linked to earlier onset of PSMD and FW by 2.0 and 2.2 years, respectively. Having two or more CMCs was consistently associated with earlier onset across all five biomarkers: 0.6 years for WMH, 2.0 years for Genu-FA, 1.4 years for PSMD, 1.6 years for FW, and 1.0 year for ARTS. Neither *APOE ε4* carriership nor level of education showed statistically significant effects on the age of onset for any biomarker, except for 4 years more education being association with 1.8 years later progression of Genu-FA.

**Table 3.** Estimated covariate effects (95% credible interval) for each outcome in years. For example, males had 2.8 years later onset of WMH abnormality in comparison to females.

| Variable | WMH | Genu-FA | PSMD | FW | ARTS |
| --- | --- | --- | --- | --- | --- |
| Male vs female(ref) | -2.8 (-3.8, -1.8) | -4.9 (-7.1, -2.8) | 2.0 (1.1, 2.9) | 2.2 (1.4, 3.1) | -9.3 (-10.2, -8.5) |
| CMC, 2 more | 0.6 (0.2, 1.0) | 2.0 (0.8, 3.2) | 1.4 (0.9, 1.9) | 1.6 (1.2, 2.0) | 1.0 (0.6, 1.5) |
| Education, 4y more | 0.2 (-0.5, 1.0) | -1.8 (-3.5, -0.2) | -0.6 (-1.3, 0.1) | -0.2 (-0.9, 0.4) | -0.3 (-1.0, 0.4) |
| *APOE* ε4+ vs ε4-(ref) | 0.9 (-0.2, 2.0) | 0.1 (-2.2, 2.6) | 0.1 (-0.9, 1.2) | 0.4 (-0.6, 1.4) | 0.9 (-0.0, 1.9) |

**Figure 3:**
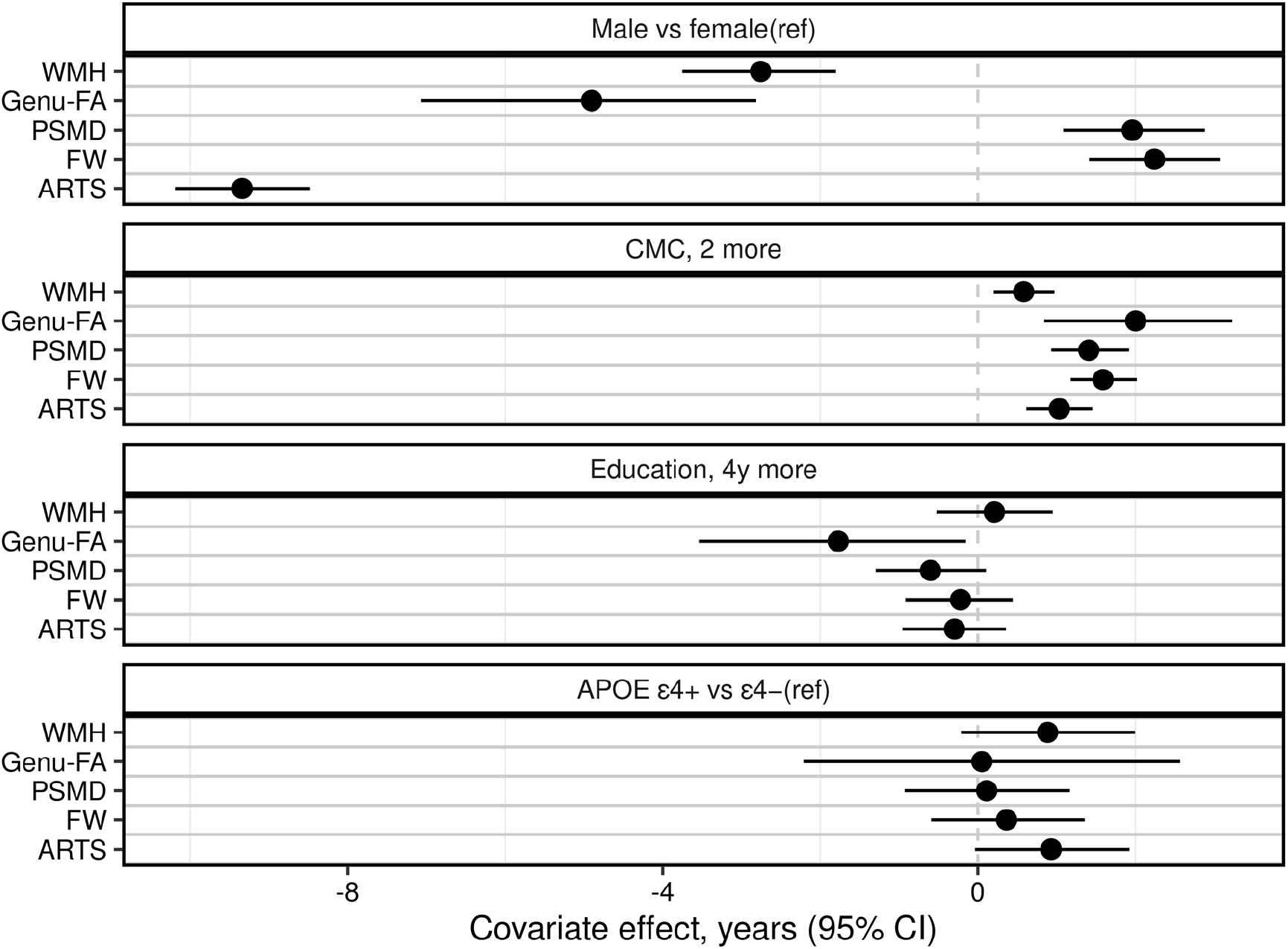
Estimated covariate effects with 95% credible intervals. A higher positive value or earlier onset relative to the population means is shown to the right of the dashed vertical line indicating that the co-variate causes an earlier abnormality of the biomarker. For example, two additional CMC was associated with earlier progression on all markers i.e. increased cardiovascular risk was associated with earlier progression of SVD biomarkers. A higher negative value or later onset relative to the population means is shown to the left of the dashed vertical line.

### Biomarker progression on the population level

**Figure 4**. displays the estimated age-related trajectories of SVD biomarker progression in the MCSA cohort, based on the proportion of participants exceeding the defined abnormality cut-points. According to the cut-point based modeling results, Genu-FA starts becoming abnormal first, followed sequentially by ARTS, FW, PSMD, and lastly, WMH. The progression curve for Genu-FA rises more gradually than the others, indicating an earlier onset but a slower rate of abnormal accumulation. In contrast, the trend of percent over threshold for the four dMRI markers was more similar. Based on the age at which we predicted 50% abnormality of each biomarker, ARTS is followed by FW with a lag of approximately six years, which is followed by PSMD by approximately four years after FW, while WMH was last and seven years after PSMD. The shallow slope of the Genu-FA curve results in crossover with the other markers, suggesting that some individuals may exhibit abnormalities in ARTS, FW, PSMD, or even WMH before meeting the criterion for Genu-FA abnormality. The age intervals between the percentage over threshold curves remain relatively constant across FW to ARTS, ARTS to PSMD, and PSMD to WMH.

**Figure 4:**
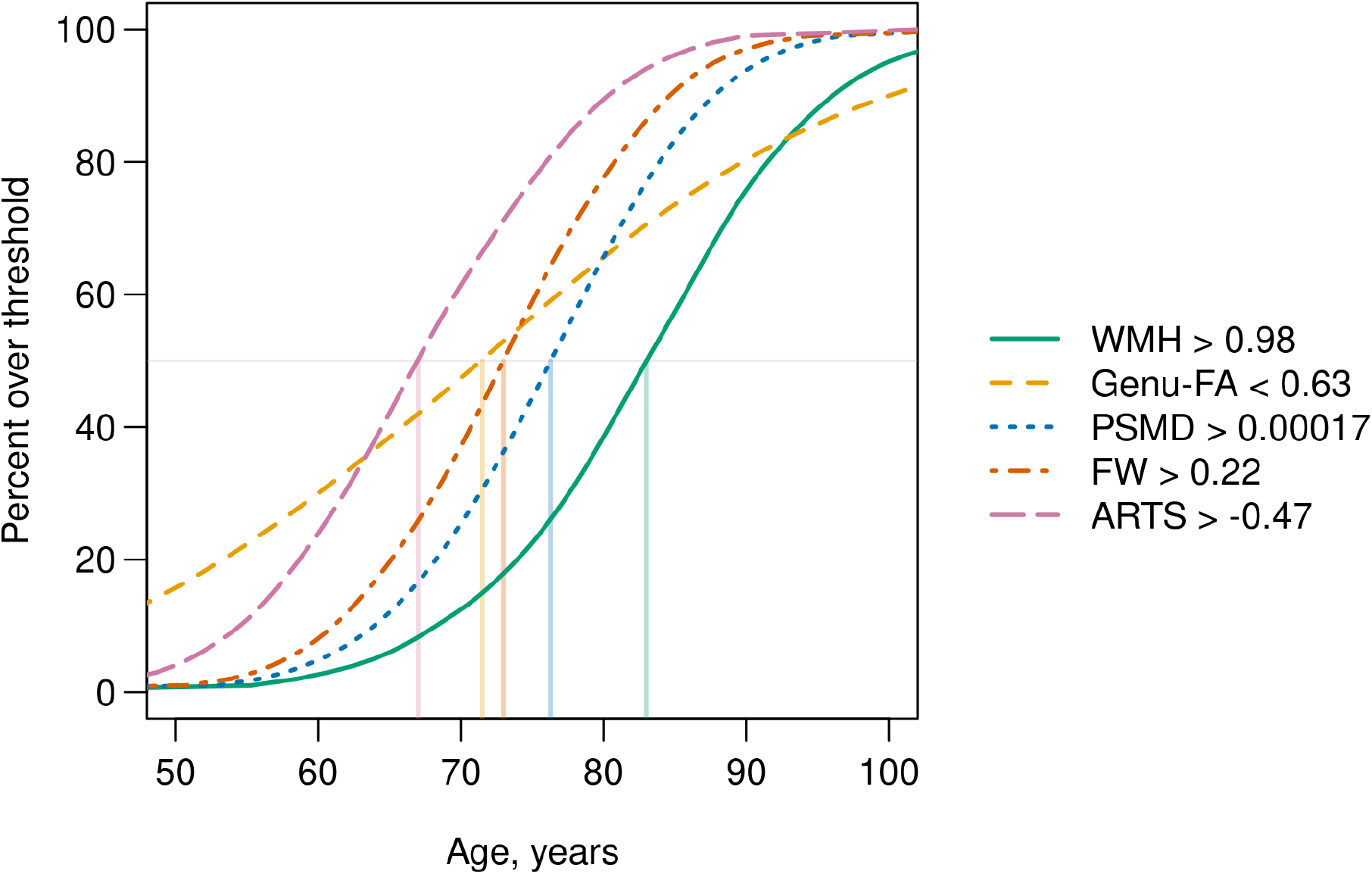
Timing of SVD biomarker progression reflects the estimated relative timing of progression of SVD biomarkers based on the fitted NLME model. The cut-points of the SVD measures that indicate abnormality of the measures are shown on the right legend. The age at the predicted value for each biomarker has also been indicated. The vertical lines indicate the ages at which 50% of participants are estimated to cross the abnormality thresholds for each of the SVD biomarkers: ARTS=67 years; Genu-FA=71.5 years; FW=73 years; PSMD=76.3 years; WMH=83 years. The units for PSMD are mm^2^/s; Genu-FA and FW are unitless; ARTS-scores represent machine learning outputs; WMH are % of the total intracranial volume.

## DISCUSSION

We evaluated the temporal progression of SVD biomarkers. Our primary findings were: 1) Early diffusion-based MRI markers precede WMH by a decade or more. ARTS, Genu-FA, FW, and PSMD became abnormal in 50% of the population approximately 16, 12, 10, and 7 years earlier than WMH, respectively. 2) There were two distinct temporal patterns of progression among the diffusion MRI based biomarkers: global markers (ARTS, FW, PSMD, WMH) were correlated and likely reflect a shared substrate of widespread WM injury; whereas regional marker (Genu-FA) demonstrated weaker correlation with WMH and had an early but gradual linear worsening across adulthood. 3) Except *APOE ε4* status, all covariates had an impact on SVD progression: sex differences in SVD progression were biomarker-specific, with females exhibiting earlier WMH, Genu-FA, and ARTS abnormalities, whereas males demonstrated earlier abnormalities in PSMD and FW. Higher CMC predicted earlier abnormalities in SVD biomarkers. Higher education showed a later Genu-FA abnormality.

### Prolonged window for early intervention and treatment of SVD

SVD is a prominent contributor to age-related cognitive decline, mortality, stroke, and gait impairment. Early SVD manifests prominently in WM because of its intrinsic vascular vulnerabilities, and it contributes to tissue injury through multiple interacting mechanisms, including blood brain barrier dysfunction, neurogliovascular unit and endothelial dysfunction, chronic hypoperfusion leading to ischemia and WM rarefaction^3^. This early tissue damage can be quantified using diffusion MRI which measures microstructural WM alterations through decreased FA and increased MD. Longitudinal diffusion MRI studies have shown that baseline microstructural damage in normal appearing WM predicts future WMH^7,24^. In the last decade, several markers have been proposed to quantify SVD-related damage from diffusion MRI that slightly vary in processing and quantification approaches, but all aim to capture WM microstructural injury^9,11,12,25^. We have included three of these diffusion MRI measures here, along with ARTS which utilized information from diffusion MRI along with WMH and demographics and found that all the diffusion MRI biomarkers change before the appearance of significant WMH. These timing curves illustrate the value of NLME models for characterizing the temporal dynamics of both conventional and emerging SVD markers. Previously WMH has been shown to precede lacunar infarcts^26^ and therefore estimating the timing of diffusion MRI is crucial for estimating the timing of early SVD. Unlike prior studies evaluating SVD biomarkers, our study quantified the timing of when these biomarkers become abnormal relative to WMH and provides evidence for a prolonged window for early diagnosis, risk stratification, and intervention in SVD highlighting the usefulness of diffusion MRI based biomarkers. Identifying SVD prior to the development of WMH may extend a potential treatment window.

### Temporal patterns across SVD biomarkers: Timing and Differences

PSMD measures the spread/dispersion of the MD within the global WM skeleton and has been shown to be particularly sensitive to vascular processes^11^. In a multi-cohort study across the lifespan, PSMD accelerated around age 70 years^27^ which is consistent with our results showing the steepest slope of PSMD abnormality increase in the 70s. Single-shell FW, originally proposed by Pasternak et. al.^28^, has been implemented differently in MarkVCID. Instead of relying on spatial regularization ^29,30^, it requires assuming a constant tissue MD resulting in high correlation between MD and FW^12^. Therefore, it was not surprising to see that PSMD and FW individual adjustments were most highly correlated (Table 2). Further, PSMD has been shown to be a strong predictor of WMH^31^. Together, the tight coupling between PSMD, FW, and WMH reflect shared substrate of widespread WM injury.

Brain arteriolosclerosis is a major contributor to SVD characterized by progressive narrowing, thickening, and stiffening of small penetrating arteries and arterioles in the brain. Currently there are no *in vivo* methods available to directly measure arteriolosclerosis and it is quantified using neuropathology. ARTS uses a machine learning model trained on neuropathology data using information from regional FA, WMH, and demographics to predict likelihood of arteriolosclerosis on new participants^14^. Therefore, moderate correlation of ARTS with WMH and other diffusion MRI markers are expected. The earlier onset of ARTS trajectories may be expected because it utilizes multimodal information including age in the model and therefore sensitive to early SVD.

Genu-FA was proposed to capture early, diffuse microstructural injury to the structurally and anatomically vulnerable frontal hemispheric connections^9^, consistent with well-established anterior-to-posterior gradient of WM vulnerability in aging and SVD^32^. There is evidence that changes to Genu-FA are seen very early with worsening of vascular risk^10^ and likely precede downstream WMH^5^. The relatively weaker linkage we observed between the timing of Genu-FA and WMH progression (than the correlation between other markers and WMH) suggests that frontal regional WM susceptibility may start early and evolve independently of overt lesion burden. Nelson et. al. mapped cerebrovascular disease as a function of age using neuropathology data in the National Alzheimer’s Coordinating Center (NACC) Registry and found a linear increase in cerebrovascular disease starting at 15% at age 50^33^. The temporal progression of Genu-FA was similar with a linear trend and >15% of participants with Genu-FA abnormalities at age 50. Our recent neuropathology study also provided evidence of a strong association between Genu-FA and global cerebrovascular neuropathology scales^34^. These data taken together suggest that Genu-FA is a useful indicator of global cerebrovascular damage and Genu-FA abnormality will inform on early SVD.

### Risk Factors of temporal progression of SVD

Worse CMC (cardiovascular metabolic conditions) consistently shifted SVD biomarker trajectories earlier, with effects observed across all imaging markers. This finding reinforces the cumulative impact of modifiable cardiovascular risk factors on both microstructural and macrostructural WM damage and supports aggressive midlife cardiovascular risk management. Education showed a selective protective association with Genu-FA. Given that Genu-FA had a comparably stronger association with CMC, the association between education and Genu-FA abnormalities may suggest that Genu-FA is capturing early SVD.

Sex differences were biomarker specific with females exhibiting earlier WMH, Genu-FA, and ARTS abnormalities, whereas males demonstrated earlier abnormalities in PSMD and FW. Females have greater burden of cerebral arteriolosclerosis and SVD, despite less large vessel atherosclerosis than men supporting the ARTS and WMH findings^35,36^. Sex differences in diffusion MRI have been reported in the literature^27,37^. FA in the anterior corpus callosum is known to be lower in females even in healthy controls^38^. Changes observed in PSMD and FW (derivatives of MD) are reflective of global WM damage, and it was recently shown in a dataset of >12 K participants that tissue loss with age was accelerated in males^39^ supporting our findings. *APOE ε4* status was not associated with the onset of SVD biomarkers, reinforcing the distinction between SVD driven white matter injury and Alzheimer’s disease pathology that is driven by *APOE ε4* status.

### Strengths and Limitations

A key strength of this work was the utilization of multi-output nonlinear mixed-effects modeling with a large, longitudinal neuroimaging dataset that enabled direct comparison of biomarker timing on a shared age axis, providing a robust framework for disentangling heterogeneous SVD trajectories and capturing individual-level variability in onset of SVD abnormalities. However, a limitation was that it assumes that all participants follow the same trajectory. While the cut-points influence the determination of abnormalities, the utilization of 30-49 year old cognitively unimpaired participants and participants with vascular cognitive impairment for determining (the cut-point for) abnormalities for each diffusion MRI based biomarker supports our findings of timing differences observed between the SVD biomarkers. Another limitation is the use of a predominantly white sample representative of the Olmstead County population that it was sampled from.

## CONCLUSION

This work supports a shift toward diffusion MRI based biomarkers for early SVD detection, with implications for identifying high-risk individual’s decades before substantial WMH burden and subsequent irreversible cognitive and functional decline are observed.

## Acknowledgements/funding sources/conflicts

### Funding

This work was supported by NIH grants UF1NS125417 (PI: Vemuri and Petersen), R01 AG056366 (PI: Vemuri), U01 AG006786 (PI: Petersen), P50 AG016574 (PI: Petersen), R01 AG034676 (PI: Rocca), R37 AG011378 (PI: Jack), R01AG054787 (PI: Windham); the GHR Foundation grant, the Alexander Family Alzheimer’s Disease Research Professorship of the Mayo Foundation, the Elsie and Marvin Dekelboum Family Foundation, U.S.A. and Opus building NIH grant C06 RR018898.

## Acknowledgments

We thank all the study participants and staff in the Mayo Clinic Study of Aging, Mayo Alzheimer’s Disease Research Center, and Aging Dementia Imaging Research laboratory at the Mayo Clinic for making this study possible. We gratefully acknowledge the support of NVIDIA Corporation for the donation of the Quadro P5000 GPU used in this research. The codes used to run Free water and ARTS pipelines for the preparation of this article were funded by NINDS/NIA as part of the MarkVCID Consortium (U24NS10059). A complete listing of MarkVCID investigators can be found on https://markvcid.partners.org/acknowledgements.

## Data availability

Mayo Clinic Study of Aging data are available for download on the GAAIN website: https://www.gaaindata.org/partner/MCSA

